# *Ostrea edulis* at shipwrecks in the Dutch North Sea

**DOI:** 10.1101/2020.01.09.883827

**Authors:** Joop W.P. Coolen, Udo Van Dongen, Floor M.F. Driessen, Erik Wurz, Joost H. Bergsma, Renate A. Olie, Brenda Deden, Oscar G. Bos

## Abstract

During diving expeditions in July and September 2019 two live European flat oysters (Ostrea edulis) were observed by SCUBA divers. The first oyster was found lying amongst coarse shelly material in the scour hole around shipwreck #3251, 37 nautical miles north west of Texel (NL). The second oyster was found on the Gustav Nachtigal wreck, 6 nautical miles north of Schiermonnikoog (NL). Additional shipwrecks were also inspected for O. edulis individuals but no live specimens were observed, although on 9 out of 11 inspected locations empty, fossil O. edulis shells were found. The September dive on the #3251 shipwreck revealed the presence of several large empty flat oyster shells, some of which were attached to each other, with up to three individuals in a cluster. This implies that the area around the #3251 shipwreck was a suitable location for flat oysters in the past and that oyster larvae are still capable of reaching the location. The findings suggest that shipwrecks and their surroundings are promising locations for future O. edulis restoration projects.

## INTRODUCTION

The European flat oyster *Ostrea edulis* once formed an extensive population in offshore Dutch waters (Olsen, 1883), that has been overfished and was decimated around 1900 (Kamermans et al., 2018). Nowadays, the flat oyster is considered extinct offshore in the Netherlands (Berghahn and Ruth, 2005; Christianen et al., 2018; Kamermans et al., 2018). Although coastal populations have been reported recently in the south of the Netherlands, near shore and offshore in Belgium and in the Dutch part of the Wadden Sea (Bos et al., 2019; Christianen et al., 2018; van der Have et al., 2017; Kerckhof et al., 2018), no living *O. edulis* of natural origin have recently been recorded in Dutch offshore waters north of the Offshore Wind Farm Egmond aan Zee (OWEZ) where the most northern recent living offshore oyster in the Netherlands was found (Bouma and Lengkeek, 2013, but see Kerckhof et al., 2018).

Here we report the most northern findings of live *O. edulis* specimens in the Dutch North Sea, on shipwrecks and their surrounding scour holes.

## MATERIALS AND METHODS

In July and September 2019 a group of SCUBA divers visited 22 wrecks in the Dutch and British part of the southern North Sea (Table 1). The SCUBA divers performed a visual biodiversity inventory for a wide range of species and collected specimens for later identification, as described in Schrieken *et al.* (2013).

**Table 1.**
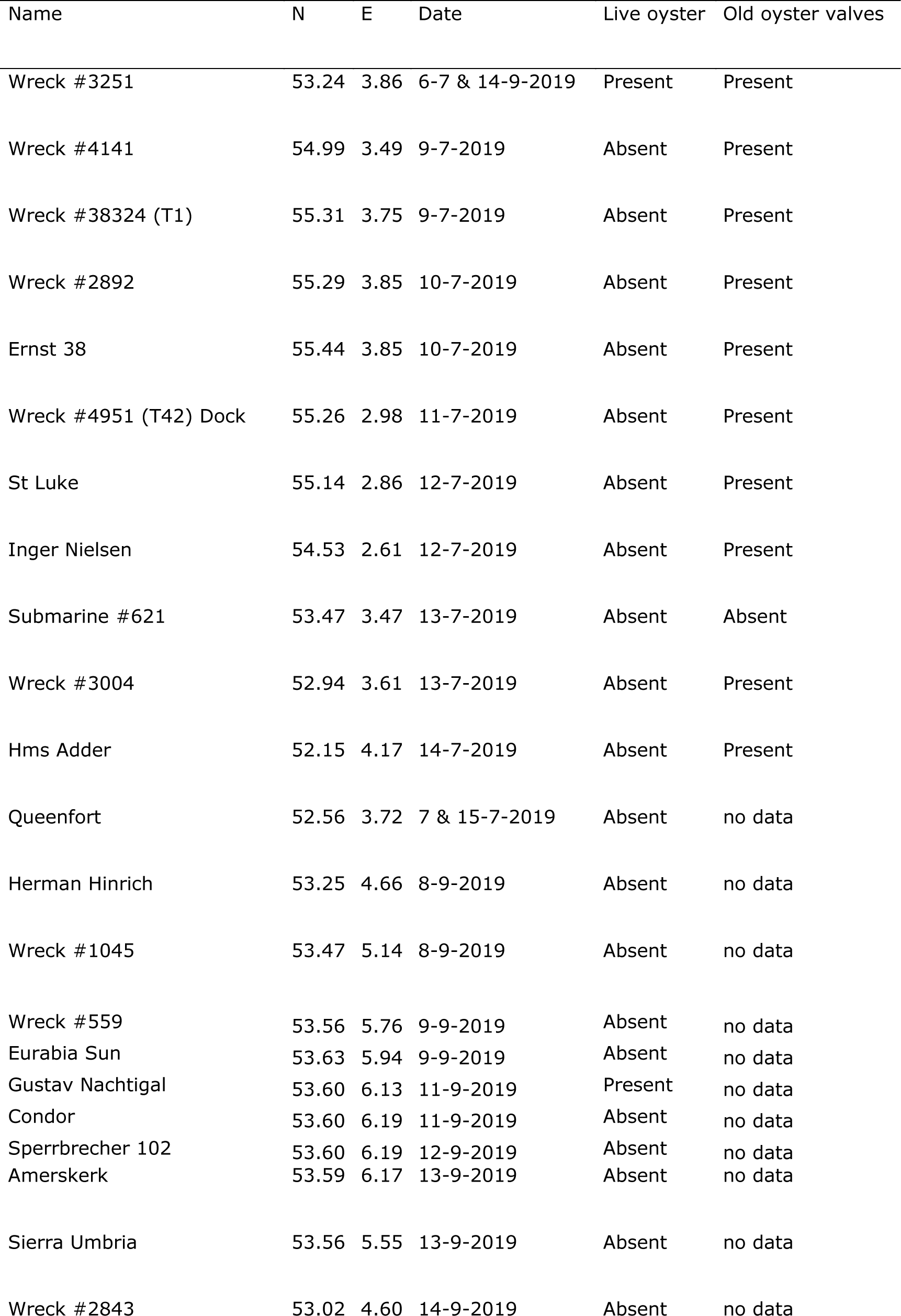
Location information. Wreck names/numbers or descriptions, positions (WGS84), survey dates, presence of live oysters or empty shells on the wrecks visited.

During these expeditions a wreck of an unknown steam ship (wreck number #3251) 37 nautical miles north west of the island of Texel (NL) was visited on 6 July and 14 September 2019. The wreck is located at 53.457°N 3.911°E (Figure 1), on a sandy bottom in a maximum water depth of 30 meters. Based on the size of the screw and rudder blade, participating archaeologists judged the wreck to be from a cargo vessel (personal communication Betty van den Berg and Robertino Mulder, Dive the North Sea Clean Foundation). Given that the vessel was a steam ship, it is likely to have been wrecked roughly 100 years ago.

**Figure 1.**
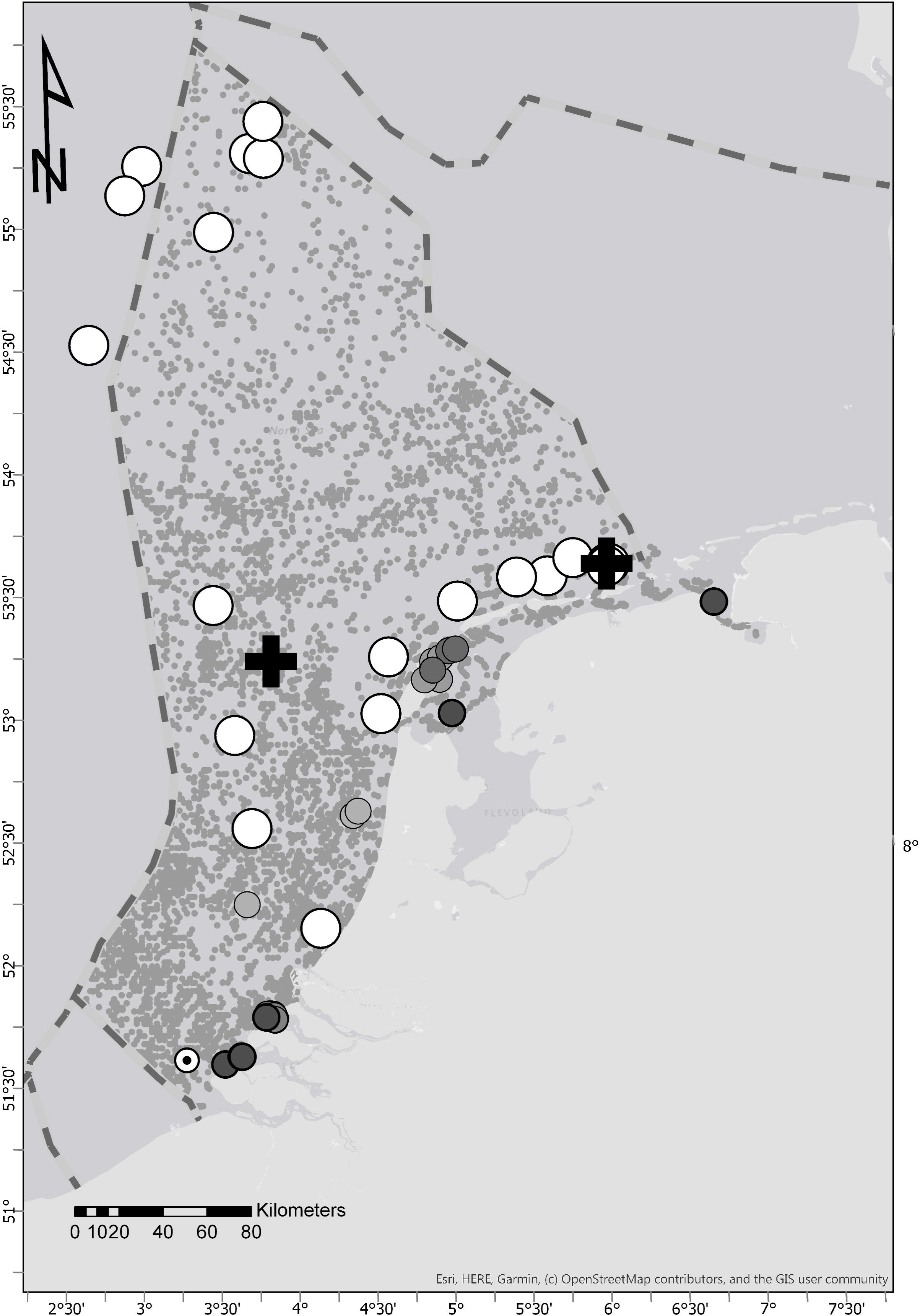
Locations of sightings of *Ostrea edulis*. Black plus signs: sightings on the #3251 and Gustav Nachtigal wrecks. Large white dots: absence observations on all other wrecks inspected in July and September 2019. Medium sized grey dots off the coast: sightings offshore (OWEZ windfarm and HMS Aboukir; Kerckhof et al., 2018). Medium sized white dot with black centre: unconfirmed sighting offshore (Christiaan Huygens wreck; Kerckhof et al. [2018]). Medium sized light grey and intermediate grey dots: sightings during dedicated one-time inventories (Bos et al., 2019; Christianen et al., 2018; van der Have et al., 2017). Dark grey medium sized dots: presence observations from fish and shell fish surveys extracted from FRISBE-database (data 2007-2017), Wageningen Marine Research. Small grey (background) dots: absence sightings from fish and shellfish surveys (Bos et al., 2019; Christianen et al., 2018; van der Have et al., 2017). Striped line: Borders of exclusive economic zones of Belgium, the Netherlands, Germany, Denmark and United Kingdom. Tick borders with degrees north and east in WGS84.

On 11 and 21 September 2019 the wreck of the Gustav Nachtigal was visited. The Gustav Nachtigal was a speedboat escort ship, attacked and sunk in 1944 and is since then located at 53.600°N 6.133°E, in 23 meters water depth, 6 nautical miles north of Schiermonnikoog.

During all dives, visited wrecks were inspected for presence of live *O. edulis* specimens. If present, specimens were collected if possible, or photographed when collection was not possible. In July, presence of empty valves of *O. edulis* around shipwrecks was registered as an indication of historic presence of *O. edulis* in the area. Furthermore, during the visit of wreck #3251 in September, empty *O. edulis* valves were collected. The length and width of a selection of the collected specimens was measured to the nearest mm.

Identification of *O. edulis* was based on a clear flattened right valve and concave left valve and round appearance, in combination with an irregular and rough surface of the shell when visible. Also, *O. edulis* has a relatively straight opening compared to *Crassostrea gigas*. In the collected specimen, presence of chomata in the hinge was confirmed, which is an important characteristic to differentiate between *O. edulis* and *C. gigas* (Amaral and Simone, 2014; Kerckhof et al., 2018).

## RESULTS

During the first dive in July 2019 on the #3251 shipwreck one live individual of *Ostrea edulis* was observed. The flat oyster was present lying flat on top of the coarse shelly seabed of the scour holes directly surrounding the shipwreck. The shells around the oyster consisted of *Neptunea* spp., *Ensis* spp., *Laevicardium* spp. and other smaller shelly material. The oyster appeared to have originally been attached to the shelly material but was now much larger than the fragment it was attached to. The original attachment fragment could not be further identified. In addition to a live specimen several empty grey single valves of much older European flat oysters were observed, which is common around many shipwrecks in the southern North Sea (personal observation by the authors), but none were collected in July. The specimen was collected for confirmation in the Wageningen Marine Research benthic laboratory and stored in ethanol 99% in the Wageningen Marine Research reference collection (number 2019IRC010030). The specimen measured 68 x 78 mm (Figure 2) and was estimated to be 3-4 years old. Since the oyster was found at the end of a dive, no search for additional oysters could be performed at that point. Every other shipwreck visited after that during the July and September expeditions was inspected for oyster presence as well. Only grey, old oyster valves were observed on most other shipwrecks visited in July (Table 1).

**Figure 2.**
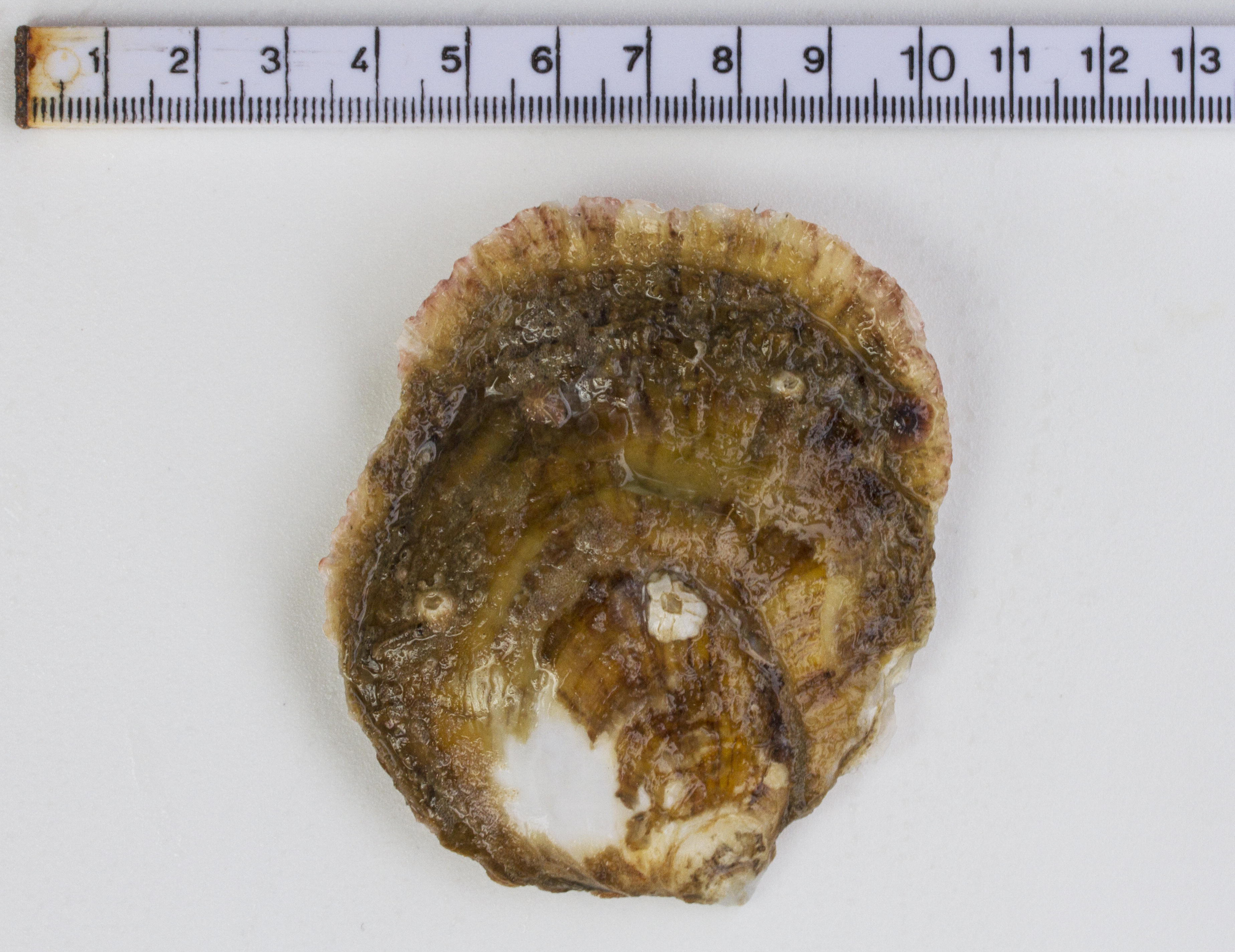
*Ostrea edulis* on #3251 wreck. Top view photograph of the live *Ostrea edulis* individual found on the seabed amongst the shelly material around the #3251 wreck in July 2019, with ruler (cm).

During a dive on the Gustav Nachtigal wreck on 11 September, a live adult flat oyster was observed (Figure 3), after which is was again observed on 21 September. The oyster was located in a sheltered area, under a small overhang. It was attached to vertical standing inside material of the wreck itself and was not surrounded by other large sessile organisms. The estimated length of the oyster was 14-18 cm, with an estimated width of 10-14 cm. The specimen was photographed *in situ* but not collected.

**Figure 3.**
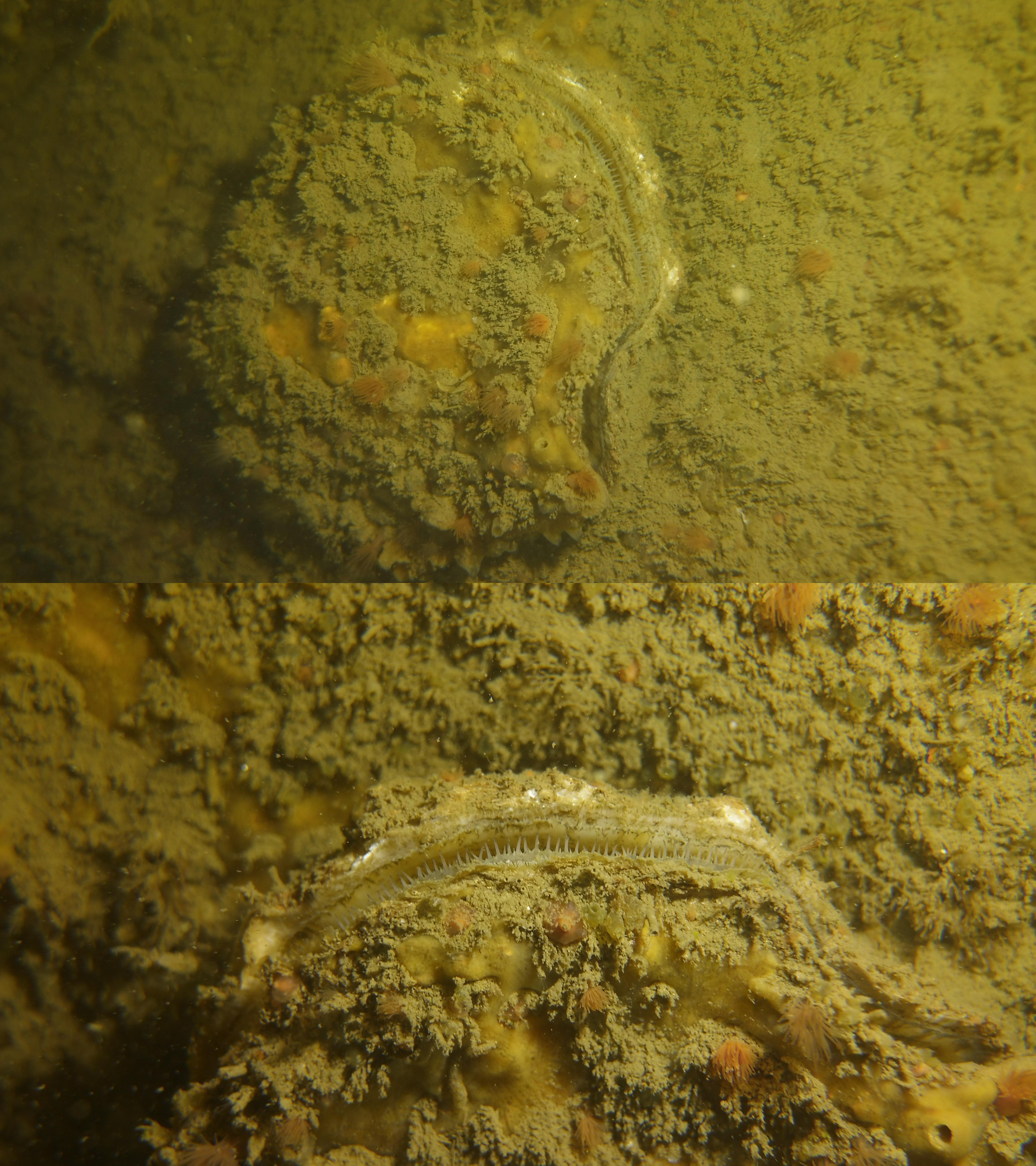
*Ostrea edulis* on Gustav Nachtigal wreck. Photographs of the live *Ostrea edulis* individual found on the shipwreck of Gustav Nachtigal in September 2019.

Grey, old oyster valves were not registered in September, except on the #3251 shipwreck. Here, several empty valves were observed and some larger specimens were opportunistically collected. Several of the observed valves were found to be growing attached to other *O. edulis* shells, with up to three valves per cluster (Figure 4). The valves were between 89 and 120 mm long and between 76 and 115 mm wide (Table 2).

**Table 2.** Measurements of empty shells from #3251 wreck. Lengths and widths in mm of a selection of the larger empty *Ostrea edulis* shells found in and around the #3251 wreck in September 2019, as shown in Figure 3.

| Cluster | Length | Width |
| --- | --- | --- |
| A1 | 116 | 112 |
| A2 | 107 | 99 |
| A3 | Small shell fragment not measured |  |
| B | 100 | 76 |
| C1 | 106 | 94 |
| C2 | 88 | 78 |
| C3 | 89 | 81 |
| D | 120 | 104 |
| E | 118 | 83 |
| F | 99 | 115 |

**Figure 4.**
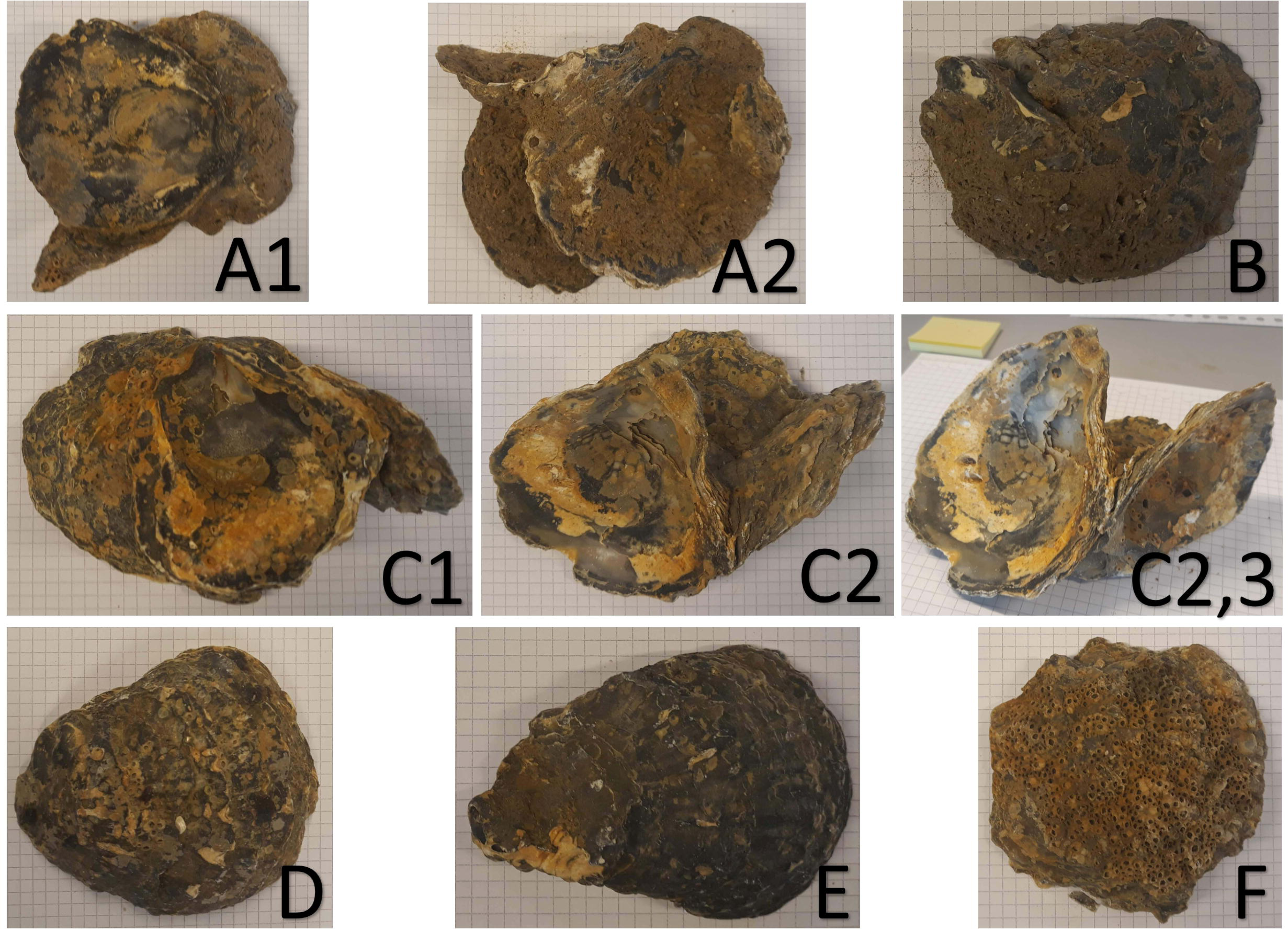
empty Ostrea edulis shells from #3251 wreck. Top view photographs of a selection of the larger empty *Ostrea edulis* shells found in and around the #3251 wreck in September 2019, with 5 mm grid, for sizes see Table 2. C2,3 with C2 on the left and C3 on the right.

## DISCUSSION

To date, the European flat oyster was considered extinct offshore in the northern area of the Dutch part of the North Sea. Only in southern Dutch North Sea waters and in the Wadden Sea flat oysters had been observed recently (Bouma and Lengkeek, 2013; Christianen et al., 2018; Kamermans et al., 2018; Kerckhof et al., 2018). During visits of over 100 ship wrecks between 2010 and 2019 by the same group of wreck divers, no other live flat oysters were observed offshore (personal observations by the authors), with exception of locations in the south of the Netherlands and Belgium (Kerckhof et al., 2018). Furthermore, recent investigations of fouling communities on oil and gas structures near shipwreck #3251, did not reveal any flat oysters (Coolen et al., 2018), nor did a study of rocky reefs in the Borkum Reef grounds near the Gustav Nachtigal wreck (Coolen et al., 2015a).

The first oyster was found amongst the coarse shelly material present in the scour pits around the ship wreck. Oysters are known to prefer clean shells as settling substrate (Brett et al., 2011). Given that shipwrecks tend to be covered in extensive marine growth (Coolen et al., 2015b; Gittenberger et al., 2013; Lengkeek et al., 2013), the shelly material was surprisingly clean. Shelly material as observed around the #3251 wreck, is very common near offshore shipwrecks in the North Sea (personal observation by authors). Therefore, these clean shells in the scour pits may be suitable material for larval settlement if the locations are within reach from a source population.

The finding of the second oyster on a shipwreck implies that wreck material, if not already fully covered by other organisms competing for settlement space, has the potential to be suitable settling substrate too. The less exposed locations of both oysters suggest that flat oysters could have (competitive) advantage of reduced current velocities. The size of the second oyster shows that there are still some older oysters present above the Dutch Wadden islands. Observations reported by Kerckhof et al. (2018) also included an oyster sheltered by wreck material, attached to steel parts of the wreck (personal observation Joop W.P. Coolen).

To assess the potential to restore offshore flat oyster populations, Kamermans et al. (2018) investigated the dispersal potential of flat oysters in Dutch waters by modelling the distribution of larvae from a set of hypothetical source locations in offshore wind farms. The general direction of dispersal of the larvae was oriented south to north along the southern Dutch coast, and west to east above the Wadden Islands. A study of *Mytilus edulis* larval dispersal in the same area predicted similar distribution paths (Coolen et al., 2019). This suggest that source locations for the observed oysters must have been south of the #3251 wreck and west of the Gustav Nachtigal. However, none of the larval dispersal scenarios explored by Kamermans et al. (2018) included larval source locations that included the location of these wrecks. Based on this we hypothesise that the source of the oysters found here must have been present at an unknown offshore location for the #3251 wreck and unknown location, possibly more near shore, for the Gustav Nachtigal wreck. Given that both oysters were several years old, they cannot have originated from the flat oysters that were introduced to the Luchterduinen wind farm, Gemini wind farm nor Borkum Reef grounds in 2018 (Sas et al., 2019). As *O. edulis* has been found in the Wadden Sea (Figure 1), this might have been the source for the Gustav Nachtigal oyster, although larvae sources on other artificial or natural reefs in the neighbourhood cannot be excluded.

The presence of a live specimen as well as empty and clustered valves, suggest that the area around the #3251 shipwreck was a suitable location for flat oysters in the past and that oyster larvae are still capable of reaching the location. The findings suggest that shipwrecks and their surroundings are promising locations for future *O. edulis* restoration projects.

## CONCLUSION

Clean shelly material present in scour pits around shipwrecks and sheltered areas on shipwreck material may be suitable for settlement of European flat oyster larvae. An adult population of reproducing flat oysters must be present in offshore Dutch North Sea waters, although their exact locations remain unknown.

## ACKNOWLEDGEMENTS

We thank the volunteers and board members of the Duik de Noordzee Schoon Foundation who organised the diving expedition. Furthermore we thank the crew of the Cdt. Fourcault (IMO 7304675; NV Seatec) and Wilhelmina (MMSI 244100334). Without their professional support at all levels, the wreck dives would have been impossible. We are grateful to the participating divers and organisations such as the Zabawas foundation, who all contributed to the funding of the diving expedition. Part of the work reported in this publication was funded through the NWO Domain Applied and Engineering Sciences under Grant 14494; the Nederlandse Aardolie Maatschappij BV, Wintershall Holding GmbH, Neptune Energy and Energiebeheer Nederland B.V. We thank Babeth van der Weide for confirming the identification of the collected *O. edulis* specimen and Jan Tjalling van der Wal for creating the plot with locations.

## Ethics approval and consent to participate

Not applicable

## Consent for publication

Not applicable

## Availability of data and materials

All data generated or analysed during this study are included in this published article. The datasets generated during and/or analysed during the current study are available in the DASSH repository, url will be provided once available, we submitted the data on 10 December 2019.

## Authors’ contributions

JWPC: designed research; performed research; analyzed data; wrote the paper

UvD: performed research, wrote the paper

FD: designed research, performed research, wrote the paper

EW: performed research, wrote the paper

RO: designed research, performed research, analyzed data, wrote the paper

BD: performed research, wrote the paper

OGB: designed research, performed research, analyzed data, wrote the paper

All authors read and approved the final manuscript.

